# Repeatable flight initiation distance is associated with nest-site characteristics in a colonial seabird

**DOI:** 10.1101/2025.10.27.684831

**Authors:** Jeffrey Carbillet, Jaanis Lodjak, Hannah Métaireau, Tuul Sepp

**Affiliations:** Institute of Ecology and Earth Sciences, University of Tartu, Tartu, 51014, Estonia

**Keywords:** flight initiation distance, colonial seabird, Common gull, behavioural ecology, life history strategies

## Abstract

Behavioural responses are among the fastest ways for wild animals to respond and adapt to environmental change. In breeding birds, nest defence behaviour is particularly important, as it directly influences reproductive success. However, the variability of this behaviour, and its links to environmental conditions and fitness, remain poorly understood. Here, we investigate the repeatability of flight initiation distance (FID) – a proxy for nest defence behaviour – in Common gull (*Larus canus*) breeding pairs within a seabird colony. We assess how consistent differences in FID between gull pairs relate to nest-site characteristics and fitness-related traits. We find that FID is strongly repeatable at the pair level (R = 0.50), indicating consistent behavioural differences among breeding pairs. Moreover, pairs with shorter FID nest at higher elevations and in areas with higher conspecific density compared to pairs with longer FID. Finally, our results show that these different behavioural responses, linked to different nest-site contexts, do not lead to clear differences in terms of fitness outcomes, suggesting that breeding pairs with marked behavioural differences could achieve similar fitness by matching the environmental conditions of their nesting site with their behavioural responses to perceived risks.

## Introduction

Behaviour is a life history trait that allows animals to respond and adapt to environmental changes. Rapid, extensive human-induced environmental changes mean that species experience novel environmental conditions, where behaviour can play a leading role in allowing them to adjust, explaining why some species are able to survive, while others struggle (Wong & Candolin 2015). Unlike morphological or physiological traits, which may require multiple generations to evolve, behavioural adaptations can occur in ecological timescales, within an individual’s lifetime, thus allowing for “decision making” and immediate responses to new challenges (Lima 2009). However, constraints on behavioural flexibility exist, arising from animal personalities and costs of flexibility (Duckworth 2010), which set limits to the ability to respond to fluctuating environments. The biology of breeding birds is among the most studied topics in evolutionary ecology and is a good model for studying the potential of animals to acclimate to changing environments. As this potential is tightly linked to fitness outcomes, breeding birds have a significant capacity to assess and respond to changes in the risk of predation to themselves and their eggs/nestlings (Lima 2009). There is, however, still little empirical evidence on how much nest defence behaviour reflects repeatable differences among individuals and/or plasticity in response to environmental conditions and internal state, limiting our ability to predict the potential for rapid adaptation (Thys et al. 2019).

Colonially nesting seabirds present a particularly interesting case for studying behavioural flexibility in the context of environmental changes. Nearly one-third of seabird species are threatened due to reduction in marine food supplies, environmental contaminants and ocean acidification, invasive species, loss of nesting habitat, and global climate change (Jones & Kress 2012). Living in large, dense colonies, these birds face many challenges, including competition for resources, predation risk (including cannibalism), and sensitivity to environmental fluctuations. However, these risks are balanced with the ability of enhanced social information use (Evans et al. 2016; Monier 2024). Despite the growing body of literature on the effects of intrinsic and external factors (e.g. social and environmental) on individual behavioural traits, this issue remains poorly understood. Understanding the amount of flexibility in behavioural traits helps to predict the ability of birds to cope with (or suffer from) rapid changes in their environment. For example, a study in lesser black-backed gulls (*Larus fuscus*) showed that after loss of breeding grounds, relocated birds still used the foraging sites of their natal colony, although this resulted in longer travelling distances when compared with the residents of their new breeding grounds (Kavelaars et al. 2020). At the same time, a study in European shags (*Phalacrocorax aristotelis*) indicated that the birds could quickly respond to an introduced invasive predator by moving to nest-sites that granted greater protection (Barros et al. 2016).

One of the most easily standardisable and less-invasive measures of antipredator behaviour is the distance at which animals move away from approaching threats (flight-initiation distance, FID, Blumstein 2003). FID has been used by behavioural ecologists to understand both the trade-offs of antipredator behaviour and sensitivity to (human) disturbance (Weston et al. 2020). It is predicted that escape decisions will be flexible and are influenced by both the costs and benefits of remaining on site (Blumstein 2003). However, some studies in birds also report consistent individual differences (repeatability) in FID (e.g. Carrete and Tella 2010; Hammer et al. 2022). Although previous studies have linked behavioural consistency and flexibility to foraging, nest defence, and fitness-related outcomes in seabirds (Patrick and Weimerskirch 2024; Regan et al. 2024; Lee et al. 2025), less is known about how repeatable escape responses during incubation covary with local nest-site characteristics and reproductive traits within colonies. We present three years of data on the repeatability of FID behaviour in a colonial seabird, the Common gull (*Larus canus).* Measured at the pair level (repeatability among gull breeding pairs), we link these data to nesting site characteristics — nest elevation and local conspecific density — as well as to fitness outcomes per pair based on hatching success, probability to have at least one chick, laying date, egg mass, clutch mass, hatchling mass, and brood mass. We hypothesise FID to be repeatable at the pair level, reflecting consistent among-pair differences in risk-related behaviour during incubation. We also hypothesise FID of breeding pairs to be associated with nest-site characteristics such as elevation and conspecific density. Last, we hypothesise that FID is not necessarily associated with consistent differences in fitness-related traits, which would be consistent with the idea that different risk-related behaviours are linked to different nest-site contexts (e.g. flooding exposure, competition) without leading to clear differences in terms of fitness.

## Material and Methods

### (a) Study site

The study took place in a colony of Common gulls located on the Kakrarahu islet in Matsalu National Park, West coast of Estonia (58°46′ N, 23°26′ E). The islet is situated in the northern temperate climate zone, in the Baltic Sea, and is mainly composed of limestone gravel with patches of dense vegetation of herbaceous plants, which become dominant during late May and early June. On average, approximately 1200 pairs of Common Gulls have bred in this colony each year over the past five years.

### (b) Field data collection

As part of a long-term capture-mark-recapture program initiated in 1962, 240 Common gull nests in Kakrarahu were monitored and behavioural measurements recorded each year between the beginning of May and the middle of June between 2023 and 2025. Each nest was marked with a wooden pole, and the GPS coordinates were recorded. Common gull breeding pairs were assigned a unique pair ID that was retained across years when the same pair was re-identified. We visually confirmed that all breeding pairs in our study reused the same nesting site and did not switch partners across years, when they appeared to be included in more than one year in our FID protocol. We collected 356 observations of 45 pairs in 2023, 247 observations of 49 pairs in 2024, and 276 observations of 146 pairs in 2025. Only 10 breeding pairs were included in two out of the three years considered, and none in all three years. The FID protocols were conducted for the first time right after the 1^st^ egg was laid and repeated only once a day, up to 9 days in a row, in the morning. This protocol was chosen to have breeding pairs at the same incubation stage, making the comparison more relevant. Each day, nests were checked for new eggs and to update egg status (e.g. missing, hatched), and each new egg was marked and weighed (+/- 0.01g). Hatching dates were recorded for each chick, and body mass at birth was also recorded (+/- 0.01g).

### (c) FID protocol

The protocol was always conducted by a single pair consisting of one observer and one walker. While the observer changed over the years, the walker remained the same, wearing the same clothes, to keep the same walking pace and silhouette. The walker, assimilated to the threat for the focal animals (Blumstein 2003), started walking towards a focal nest after ensuring that one parent was incubating, facing and with an unobstructed view of the walker. Before starting, the starting distance (distance between the nesting site and the walker) was recorded with a range finder. Starting distance was constrained by field conditions, with 15 meters set as a lower limit after preliminary tests, to avoid birds being already in an alert state when initiating the protocol. As soon as the nesting bird flew away, the observer yelled STOP, and the walker stopped, measuring the FID distance (distance between the point he stopped and the nesting site). Because the incubating parent could not always be identified, the FID protocol was applied at the breeding-pair level rather than the individual level. FID distances were rounded to the nearest meter.

### (d) Elevation, density, and area estimates

To assess the elevation above sea level of each nesting site, we used elevation data from the Republic of Estonia Land and Spatial Development Board, available at https://geoportaal.maaamet.ee/. Using QGIS (version 3.40.2), we attributed to each GPS location of each nesting site the corresponding elevation (in meters). In addition, for each year, we created buffers with a 5-meter radius around each Common gull nest and calculated the number of other Common nests in this buffer. This was further used to assess the density of conspecifics around each nest. Finally, we distinguished two areas on our study site, one subjected to more regular contact with humans due to the setting of the field base that has been in the same place for the past 50 years (northern part of the island), compared to the rest of the island, which has always been in contact with humans only during the nest checking process. This was to account for the potential for birds to have a different degree of habituation to humans prior to FID protocol. The spatial variation of elevation, density of conspecifics, and the two areas within our study colony can be visualised in figure 1.

**Figure 1.**
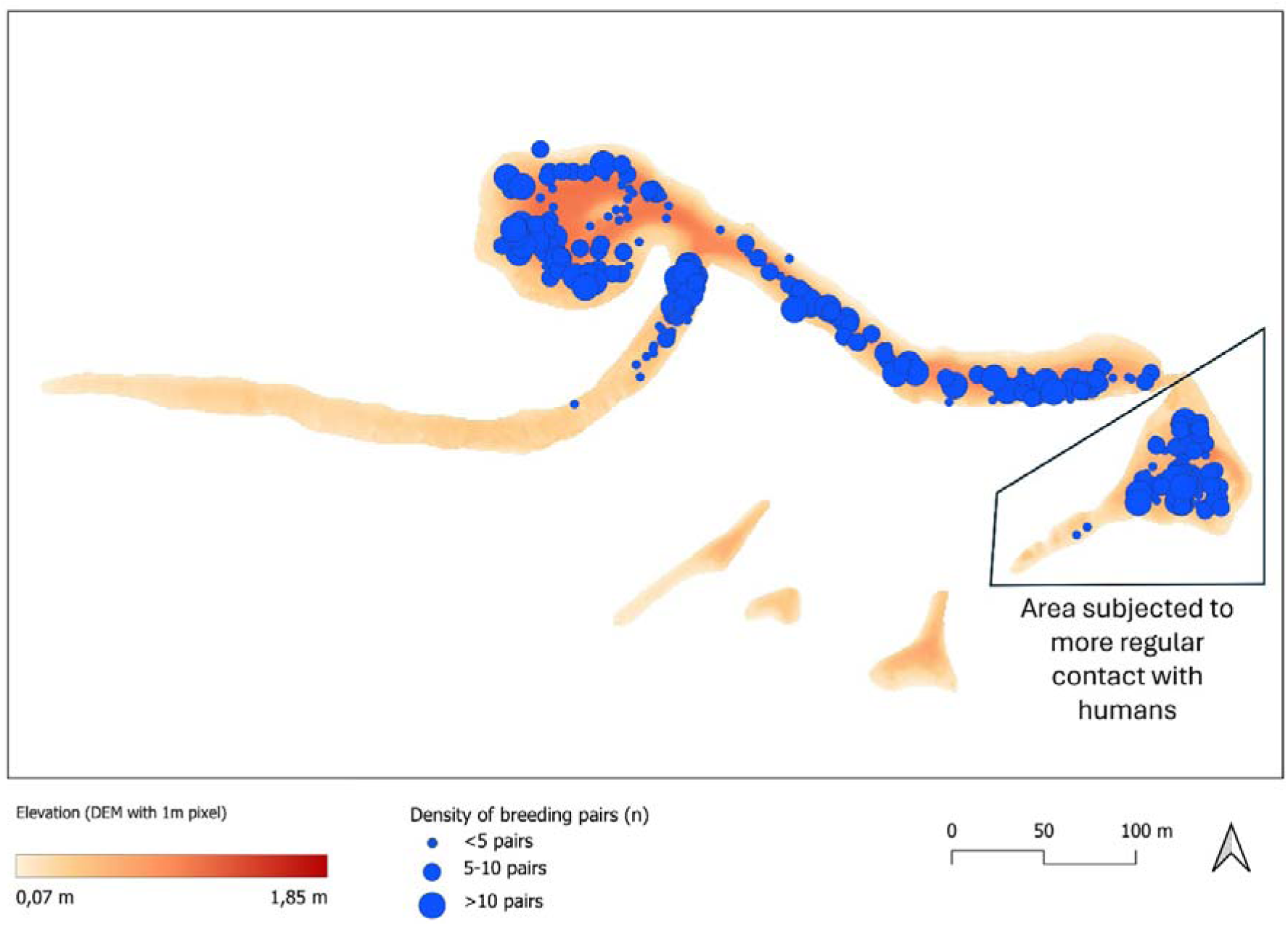
Map of the Kakrarahu Common gull breeding colony. The location of the blue dots corresponds to the nest-site of each breeding pair for which a FID protocol was conducted. The size of the dots reflects the number of conspecific breeding pairs within a 5-meter radius: small dot (< 5), medium dot (5-10), and large dot (>10). The heatmap indicates the different elevations all over the breeding colony, and the black-lined polygon highlights the area with more regular contact with humans.

### (e) Statistical analysis

Nest-level repeatability of FIDs was estimated using 879 observations of 230 breeding pairs and mixed models using the restricted maximum likelihood (REML) method, with the “rptR” package (Stoffel et al. 2017) for Gaussian distributions. The number of FID observations per pair ranged from 1 to 9, with a median of 2 and a mean of 3.8. The distribution was: 1 observation: 19 pairs; 2 observations:116 pairs; 3 observations: 2 pairs; 5 observations: 35 pairs; 6 observations: 5 pairs; 7 observations: 10 pairs; 8 observations: 40 pairs; 9 observations: 3 pairs. The variation among pairs is mostly due to early breeding failure, which prevented us from conducting more FID trials. Breeding pairs sampled only once were included in these analyses, as they still contribute to estimating the population mean, fixed effects, residual variance, and among-pair variance in the mixed model, whereas excluding them could reduce power and bias variance estimates if they were not a random subset of the population (Martin et al. 2011). Because only 10 pairs were sampled in two years, our repeatability estimate mainly reflects consistency among pairs within breeding seasons, with limited but explicit incorporation of repeated pair identities across years. The strength of the FID behaviour was assessed based on the distribution of repeatability values synthesised by Bell et al. (2009), such that values < 0.2 were considered weak, values ≥ 0.2, but ≤ 0.40 moderate, and values > 0.4 were considered strong repeatability. We calculated both the unadjusted (total variance) and adjusted repeatability for the starting distance of FID, repetition number of the FID protocol (number of observations), and the year the FID protocol trial was conducted, thereby accounting for methodological variation, seasonal timing, annual differences, and potential habituation across repeated approaches.

To analyse variation in FIDs, we performed linear mixed-effects models (LMMs) on the same 879 observations of 230 breeding pairs. These models were fitted using maximum likelihood to allow for model selection using AICc. FIDs were analysed as the response variable, and we built a set of 3 candidate models: (1) baseline, (2) link with nest-site characteristics, (3) link with nest-site characteristics in interaction with each other. For model (1), only methodological parameters were considered, such as the number of FID repetitions, the starting distance of the FID, and the area more used by humans. Model (2) added elevation of the nesting site and density of Common gull nests within a 5-meter radius, while model (3) considered their two-way interaction as well. See electronic supplementary material S1 for model details and characteristics. Pair ID and year were included as random intercepts to account for the non-independence of data and avoid pseudo-replication issues, and to control for unexplained variance due to inter-annual variation.

As a single FID behaviour value was needed for the fitness-related trait analyses, we calculated an adjusted mean FID score per breeding pair. This score was calculated as the mean residual FID per pair from a model correcting for methodological and temporal effects, including starting distance, FID repetition number, year, and area more used by humans. This allowed us to represent each breeding pair’s behavioural response with a single FID value, which we then relate to reproductive success.

For fitness-related traits, we focused on seven traits reflecting the phenology, egg and chick stages. Clutch mass, brood mass, and laying date of first egg were all analysed using linear models, while egg mass and hatchling mass were analysed using mixed-linear models, using Pair ID and year as random effects to account for the fact that several eggs and chicks originated from the same breeding pair. Lastly, the hatching success and the probability of having at least one chick were analysed using generalised linear models, with binomial distributions. The same logic using sets of 3 candidate models, as previously described, was used for each fitness trait. The main difference here was that model (2) also included mean adjusted FID as a fixed effect, while model (3) included the two-way interactions between FID and density, and FID and elevation, with no interactions between density and elevation. Details on the models for each fitness-related trait are presented in the electronic supplementary material S1.

Because FID-related predictors were tested across several fitness-related traits, we accounted for multiple comparisons using the Benjamini–Hochberg false-discovery-rate procedure. We corrected p-values across all FID-related terms tested in the fitness analyses. We report both p-values and interpret effects that were significant only before corrections cautiously.

To evaluate relative model support, we used a model selection procedure based on the second-order Akaike Information Criterion (AICc, Burnham and Anderson, 2002). Models with a difference in AICc (ΔAICc) > 2 units from the lowest AICc model were considered to have less support, following Burnham and Anderson (2002), and consequently not retained for further inference (electronic supplementary material S1). Following the approach described by Burnham and Anderson (2002) and Richards et al. (2011), we then applied a conditional model averaging procedure to estimate parameters when more than one model remained in the set of best supported models to account for model selection uncertainty. Goodness-of-fit was assessed by calculating R2, or conditional (i.e. total variance explained by the best supported model) and marginal (i.e. variance explained by fixed effects alone) R2 values (in the case of mixed models) and standard residual plot techniques (Nakagawa and Schielzeth, 2013). All analyses were carried out with R version 4.4.3, using the lmer function from the lme4 package (Bates et al., 2014).

## Results

### (a) Repeatability of FID behaviour

FID behaviour at the pair level is strongly repeatable, *R* = 0.52, 95% confidence interval = [0.45, 0.58]. When adjusted, we still obtain a strong repeatability of FID behaviour at the nest level, *R* = 0.50, 95% confidence interval = [0.43, 0.57].

### (b) FID behaviour and nesting site characteristics

In our dataset (n = 879), FIDs ranged from 0 to 67 meters, with a median value of 13 meters. Precisely, in 2023, FID (n = 356) ranged from 1 to 57 meters, with a median of 15 meters; in 2024, FID (n = 247) ranged from 0 to 57 meters, with a median of 13 meters; and in 2025, FID (n = 276) ranged from 0 to 67 meters, with a median of 10 meters. The best supported models to explain the FID behaviour variations reveal that FID is negatively associated with the elevation of the nest-site, such that breeding pairs with shorter FID are nesting at a higher elevation (variation in nest elevation in our colony is 0.28-1.76 meters), compared to breeding pairs with longer FID (table 1; figure 2). Similarly, we observe a negative relationship between FID and the density of conspecifics, such that breeding pairs with shorter FID have, on average, a higher density of conspecific pairs within a 5-meter radius of their own nest, compared to breeding pairs with longer FID (table 1; figure 3). In addition, our results show that FID decreases when the number of FID protocol trials increases (table 1) and the starting distance of the protocol decreases (table 1). Lastly, we observe breeding pairs to exhibit shorter FID, on average, in the area subjected to more regular contact with humans compared to the rest of the colony (table 1).

**Figure 2.**
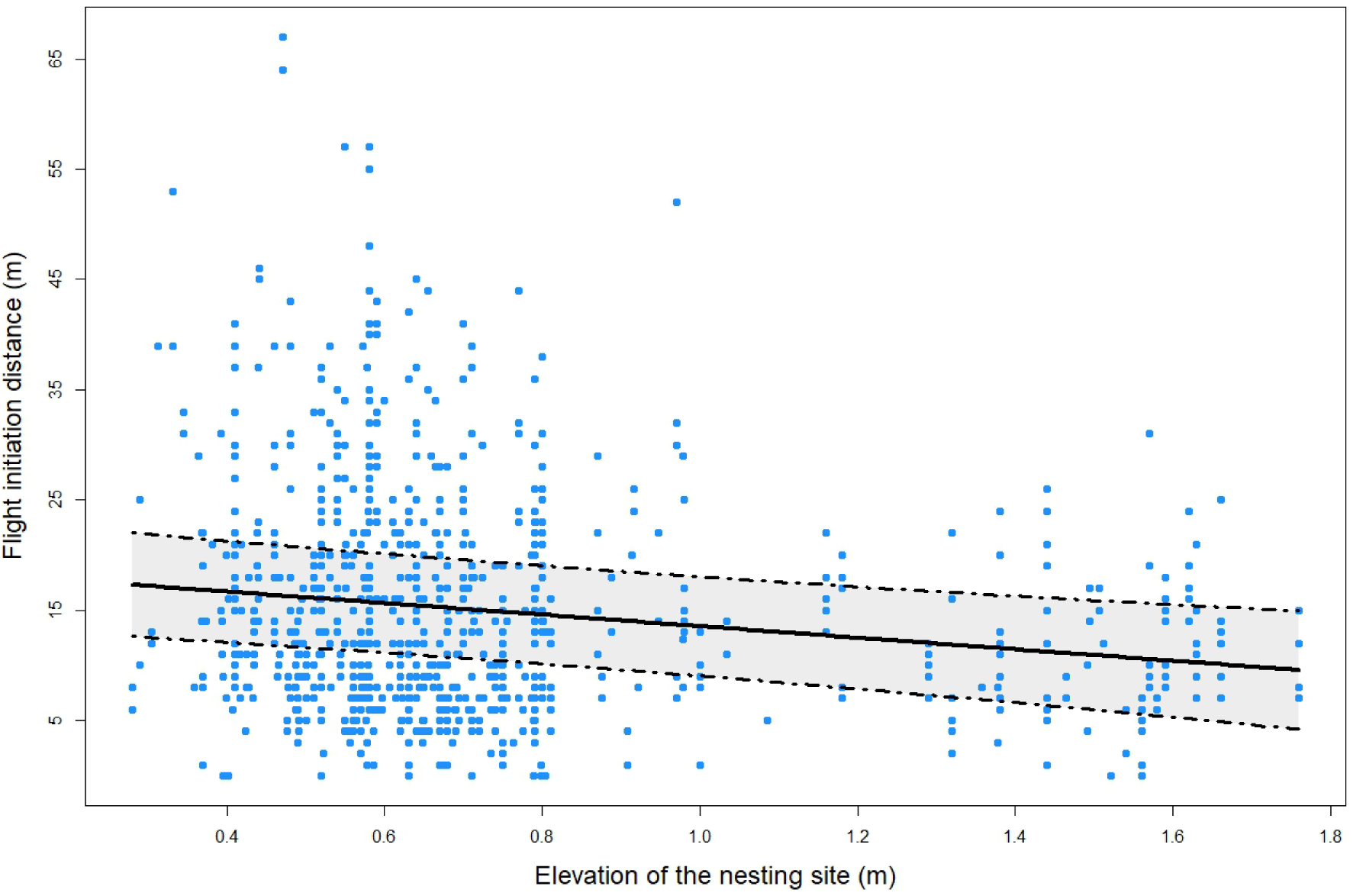
Correlation between flight initiation distances of Common gull breeding pairs and the elevation of the nesting site. Points represent observed values, solid lines the model predictions, and coloured areas in between the dashed lines the 95% confidence interval.

**Figure 3.**
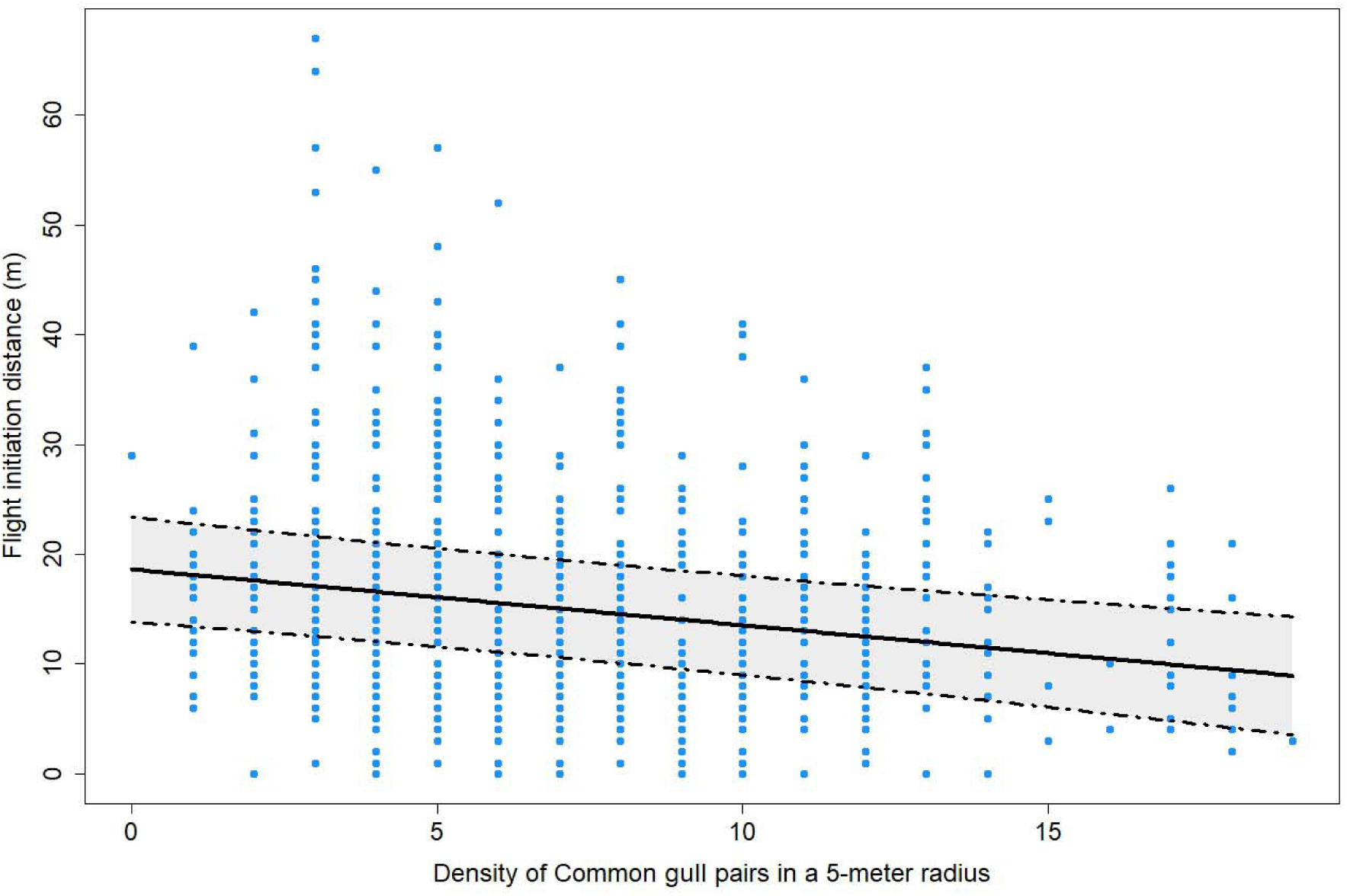
Correlation between flight initiation distances of Common gull breeding pairs and the density of Common gull pairs within 5 meters of the focal nest. Points represent observed values, solid lines the model predictions, and coloured areas in between the dashed lines the 95% confidence interval.

**Table 1.** Characteristics of the selected linear mixed-effect models for explaining variation in flight initiation distance (FID) in the Common gull breeding colony of Kakrarahu.

| Parameter | Estimate | SE | Z-value | P-value |
| --- | --- | --- | --- | --- |
| ($R^{2m}$:0.26 ; $R^{2c}$:0.64) | | | | |
| Intercept | 15.5 | 3.41 | 4.54 |  |
| Density of conspecifics | -0.60 | 0.25 | 2.38 | < 0.02 |
| Elevation of the nesting site | -6.37 | 2.33 | 2.73 | < 0.007 |
| Density*Elevation | 0.35 | 0.32 | 1.10 | 0.27 |
| Number of FID protocol trials | -0.88 | 0.14 | 6.44 | < 0.001 |
| Starting distance of the FID | 0.27 | 0.03 | 9.64 | < 0.001 |
| Area (with more frequent contact with humans) | -2.31 | 1.17 | 1.97 | < 0.05 |

$R^{2m}$ and $R^{2c}$ are the marginal and conditional explained variance. SE stands for Standard Error. See the Materials and Methods section and electronic supplementary material S1 for the definition of model sets.

### (c) Fitness-related traits and behavioural profiles

Egg mass, clutch mass, number of eggs hatched, probability of having at least one chick, brood mass, and hatchling mass showed no evidence of association with FID of the breeding pairs (table 2). On the opposite, our data suggest that when the density of conspecific pairs within a 5-meter radius increases, breeding pairs with longer FID are laying their first egg earlier than the breeding pairs with shorter FID (table 2, figure 4). However, this effect did not remain significant after correction across all fitness-related traits. We therefore interpret this pattern cautiously, as biologically suggestive rather than as strong evidence for a robust association between FID and laying date.

**Figure 4.**
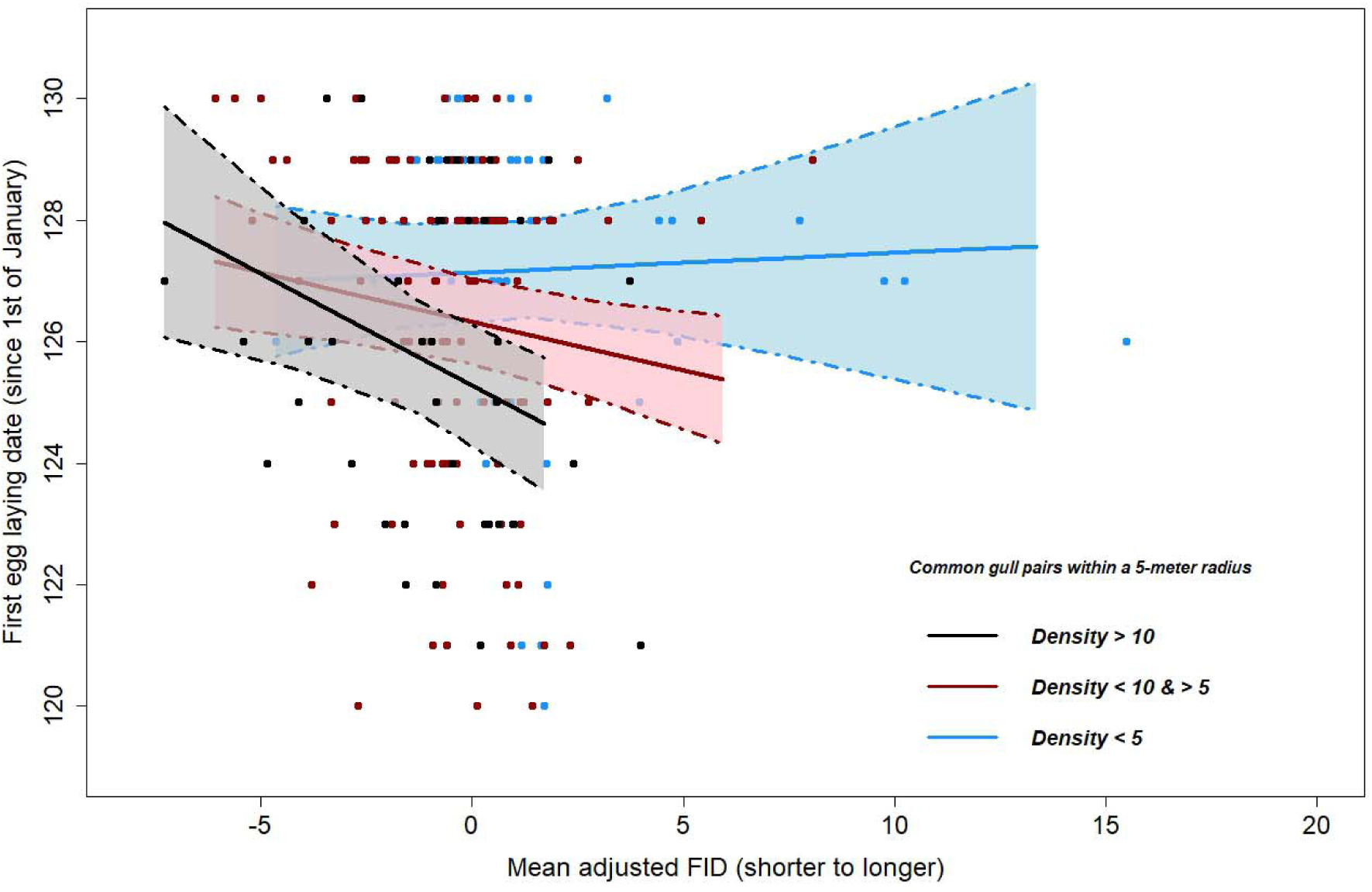
Correlation between the laying date of the first egg of Common gull breeding pairs and the adjusted mean of flight initiation distance behaviour, for breeding pairs surrounded by different densities of conspecific breeding pairs in a 5-meter radius. Points represent observed values, solid lines the model predictions, and coloured areas in between the dashed lines the 95% confidence interval

**Table 2.** Characteristics of the selected models for explaining the different fitness-related traits in the Common gull breeding colony of Kakrarahu.

| Trait | Parameter | Estimate | SE | P-value | P-BH |
| --- | --- | --- | --- | --- | --- |
| **Hatching success**<br>(n = 237)<br>Pseudo R<sup>2</sup>:0.04 | Intercept | 1.58 | 0.39 |  |  |
|  | Elevation | -0.0008 | 0.32 | 0.99 |  |
|  | Density | 0.009 | 0.03 | 0.73 |  |
|  | Year 2024 | -0.43 | 0.33 | 0.30 |  |
|  | Year 2025 | -0.97 | 0.28 | < 0.001 |  |
|  | Mean FID | -0.04 | 0.03 | 0.18 | 0.44 |
| **Probability to have at least 1 chick**<br>(n=237) Pseudo R<sup>2</sup>:0.05 | Intercept | 2.78 | 0.78 |  |  |
|  | Elevation | -0.51 | 0.55 | 0.36 |  |
|  | Density | 0.03 | 0.05 | 0.47 |  |
|  | Year 2024 | -0.71 | 0.74 | 0.34 |  |
|  | Year 2025 | -1.67 | 0.64 | < 0.01 |  |
| **Clutch mass**<br>(n=239) R<sup>2</sup>:0.89 | Intercept | 51.5 | 4.03 |  |  |
|  | Elevation | 0.12 | 2.38 | 0.96 |  |
|  | Density | 0.02 | 0.20 | 0.93 |  |
|  | Year 2024 | 4.50 | 2.28 | < 0.05 |  |
|  | Year 2025 | 3.26 | 1.90 | 0.09 |  |
|  | Mean FID | -0.14 | 0.28 | 0.61 | 0.61 |
|  | 2 eggs | 54.0 | 3.71 | < 0.001 |  |
|  | 3 eggs | 109 | 3.25 | < 0.001 |  |
|  | 4 eggs | 184 | 11.4 | < 0.001 |  |
| **Brood mass**<br>(n = 186) R<sup>2</sup>:0.78 | Intercept | 40.1 | 4.72 |  |  |
|  | Elevation | -0.24 | 3.56 | 0.95 |  |
|  | Density | 0.05 | 0.27 | 0.87 |  |
|  | Year 2024 | -0.46 | 2.88 | 0.87 |  |
|  | Year 2025 | -5.66 | 2.43 | < 0.03 |  |
|  | Mean FID | -0.30 | 0.41 | 0.48 | 0.60 |
|  | 2 hatchlings | 38.8 | 3.60 | < 0.001 |  |
|  | 3 hatchlings | 74.0 | 3.41 | < 0.001 |  |
| **1<sup>st</sup> egg laying date**<br>(n = 239) R<sup>2</sup>:0.20 | Intercept | 129.7 | 0.65 |  |  |
|  | Elevation | -0.13 | 0.58 | 0.83 |  |
|  | Density | -0.17 | 0.05 | < 0.001 |  |
|  | Year 2024 | -2.00 | 0.52 | < 0.001 |  |
|  | Year 2025 | -2.56 | 0.44 | < 0.001 |  |
|  | Mean FID | -0.11 | 0.14 | 0.41 | 0.60 |
|  | Mean FID*Elevation | 0.36 | 0.29 | 0.22 | 0.44 |
|  | Mean FID*Density | -0.4 | 0.02 | < 0.03 | 0.20 |
| **Egg mass**<br>(n = 658)<br>R<sup>2m</sup>:0.10 ; R<sup>2c</sup>:0.78 | Intercept | 55.9 | 0.84 |  |  |
|  | Elevation | -0.41 | 0.85 | 0.63 |  |
|  | Density | 0.01 | 0.07 | 0.90 |  |
|  | Egg 2 | -0.44 | 0.20 | < 0.03 |  |
|  | Egg 3 | -3.26 | 0.21 | < 0.001 |  |
|  | Mean FID | -0.06 | 0.10 | 0.54 | 0.60 |
| Hatchling mass<br>(n = 451)<br>$R^{2m}$:0.09 ; $R^{2c}$:0.50 | Intercept | 39.5 | 0.76 | | |
|  | Elevation | -0.56 | 0.80 | 0.49 |  |
|  | Density | -0.03 | 0.06 | 0.66 |  |
|  | Hatched from egg 2 | 0.31 | 0.30 | 0.30 |  |
|  | Hatched from egg 3 | -1.84 | 0.31 | < 0.001 |  |
|  | Mean FID | -0.37 | 0.30 | 0.22 | 0.44 |
|  | Mean FID*Elevation | 0.26 | 0.42 | 0.53 | 0.60 |
|  | Mean FID*Density | 0.04 | 0.03 | 0.08 | 0.40 |

N stands for the number of observations. $R^{2m}$ and $R^{2c}$ are the marginal and conditional explained variance. $P-BH$ stands for the p-value after applying the Benjamini-Hochberg correction for multiple tests. See the Methods section for more details. Pseudo $R^2$ are reported for generalised linear models, using McFadden's pseudo- $R^2$ ( $1-(\text{Residual deviance}/\text{Null}$ $\text{deviance})$ ). SE stands for Standard Error. See electronic supplementary material S1 for the definition of model sets.

## Discussion

While behavioural variability can play a crucial role in helping seabirds acclimate to rapidly changing environmental conditions, not much is known about the amount of variability in their behavioural phenotypes, and even less about how this variability is linked to life-history decisions such as nesting site selection and fitness outcomes. We report the results of a 3-year study on a standardised behavioural trait, flight initiation distance, reflecting, in most interpretations, risk-taking or nest defence behaviour, or response to human disturbance. We show that, at the breeding-pair level, FID is a highly repeatable trait (sensu Bell et al. 2009) among Common gull pairs, but is also subject to habituation effects. In our colony, seabirds that allow an observer to move closer to their nest before taking off have nests at higher elevations and have more conspecifics nesting near them. FID, in our dataset, is mostly not linked with fitness parameters.

We measured FID at the breeding-pair level, meaning that we could not (mostly) distinguish whether it was the female or male responding to our approach. In some instances, we could confirm the identity of the bird, and then we could also sometimes observe marked differences in FID between partners. The latter point indicates that individual-level variation may exist within pairs. However, the nesting site characteristics and fitness-related traits analysed here are shared by both members of the pair during a given breeding attempt. Pair-level FID therefore provides an integrated measure of the behavioural response in the context of nest defence, even though it does not allow to disentangle the separate contributions of each partner. While noise can occur due to the breeding-pair level measurement, this did not hide the high repeatability of FID (for a behavioural parameter). In previous studies, FID has been shown to be moderately repeatable in breeding Common eiders (*Somateria mollissima*, *R* = 0.4, Mohring et al. 2022), King penguins (*Aptenodytes patagonicus*, *R* = 0.26, Hammer et al. 2022), and highly repeatable in Burrowing owls (*Athene cunicularia, R* = 0.85–0.96, Carrete et al. 2016). The relatively high repeatability of FID in our study suggests that for Common gulls, FID can be used to assess relationships between behavioural response to perceived risks and other life-history traits.

The finding that breeding pairs nesting at higher locations allowed an observer to come closer before taking flight could be explained in two ways, according to the ongoing debate on whether individuals adjust their behaviour according to their local environment, or settle in habitats providing the best match to their behavioural profile (Mohring et al. 2022). First, it is possible that higher locations offer a better overview of the surroundings, and birds can allow for postponing the response behaviour. Previous studies have indicated that the environment can indeed affect FID values in nesting birds, for example, nesting in areas with denser vegetation covering the nests (Lomas et al. 2014). There are also, however, several studies finding little difference in FID values in relation to (micro)habitat selection (reviewed in Ekanayake et al. 2022). Second, birds may choose their nest locations to fit their behavioural phenotypes, with “bolder” birds choosing to nest at higher elevations (*personality matching*, Holtmann et al. 2017). For seabirds, nest site elevation is a parameter of habitat quality, as nests in lower elevations are more likely to get flooded. This also fits with the finding that nest density is related to FID, as less risk-taking pairs might find less dense areas more to their liking. With a lower number of conspecifics around one’s nest also comes lower competition to acquire and defend the nesting territory. Less risk-taking pairs may therefore trade off the advantage of nesting at a better-quality site against the cost of having to fight for it. Conversely, pairs nesting in denser areas may tolerate closer approaches because neighbouring pairs provide alarm information, reducing the perceived risk of remaining at the nest. Our correlational data cannot distinguish between these alternatives, and both processes may contribute to the observed association.

The idea that Common gull breeding pairs might match their behavioural responses with nest site characteristics is supported by the finding that fitness parameters were not statistically linked with FID. Whether it reflects predator vigilance or response to human disturbance, this link could be suspected, as theoretical models predict that the “stay or go” decision is made based (partly) on the current reproductive value (Dowling and Bonier 2018). If, however, FID is a part of a broader behavioural syndrome, and the phenotype is matched with nest site characteristics, fitness differences between less risk-taking (longer FID) and more risk-taking (shorter FID) pairs could be evened out by the choice of nest-site location. While not fully conclusive, we did find that less risk-taking breeding pairs nesting in more dense areas may lay their first egg earlier than more risk-taking pairs. This could suggest that higher density, which is related to higher competition to acquire and defend the territory in seabirds (Ellis and Good 2006), may act as a stronger selection pressure for less risk-taking breeding pairs compared to more risk-taking ones.

However, despite clear repeatable differences in FID and associations between FID and nest-site characteristics, we found little evidence that FID was strongly associated with fitness-related traits. This suggests that breeding pairs with different risk-taking behaviours occupy different nest-site contexts that may partly compensate for behavioural differences. For example, pairs with shorter FID tended to nest at higher sites and in denser neighbourhoods, whereas pairs with longer FID occupied different local conditions. Such behavioural–environment matching could reduce direct fitness differences among pairs with different FID responses. However, because our study is correlational and did not quantify nest-site availability or experimental settlement choices, this interpretation should be viewed as a hypothesis for future work rather than as direct evidence of active matching.

In our study colony, birds have been exposed to human presence for a long time, as field work has been conducted there regularly since the 1980s. Habituation can explain the pattern for lower FID values for pairs breeding in the area that is closer to our boat landing site, and where the base camp is set. In a study in female Common eiders, where birds have been similarly exposed to constant monitoring during breeding season for decades, population-level habituation was not recorded, but inter-individual variation in adjustment of FID to repeated human approaches was seen, with some bird shortening their FID over repeated trials, but others not (Mohring et al. 2025). In our dataset, within-season habituation is more marked, as FID decreases when the number of FID protocol trials increases. It is interesting to note, however, that even in these conditions of constant non-threatening exposure, habituation to FID protocol can still be seen at the breeding-pair level, indicating that birds differentiate between our everyday comings and goings, and targeted movements towards their own individual nest, with learning being involved in adjusting their response to this type of human behaviour. This indicates that even with our regular presence on the islet, there is still room for learning and habituation in the way that the gulls react towards human presence.

In conclusion, our study provides one of the rare empirical supports for consistent breeding-pair differences in behavioural responses to perceived risks, and for a link between these behaviours and environmental factors characterising nesting sites. While our results suggest that breeding pairs with marked behavioural differences could achieve similar fitness by matching the environmental conditions of their nesting site with their behavioural responses to perceived risks, further research is needed to explore this relationship in more detail. Such work is crucial for understanding the potential of wild species to rapidly adapt to environmental change through behavioural mechanisms.

## Ethics

All the procedures used in this study were approved by the Ministry of Rural Affairs of the Republic of Estonia (licence no. 1-3/25/93 and 1-3/24/115, issued 11.04.2025) and were performed following relevant Estonian and European guidelines and regulations.

## Data accessibility

Datasets and code for processing and analysing data are available at: https://doi.org/10.5281/zenodo.21158563.

## Authors’ contributions

J.C.: conceptualisation, investigation, data curation, formal analysis, funding acquisition, methodology, data curation, project administration, writing-original draft, and writing-review and editing; J.L.: investigation, writing-review and editing; H.M.: investigation, methodology, writing-review and editing; T.S.: investigation, writing-original draft, and writing-review and editing. All authors gave final approval for publication and agreed to be held accountable for the work described in this manuscript.

## Supporting information

electronic supplementary material S1

