## Supplementary material for "Repeatable flight initiation distance is associated with nest-site characteristics in a colonial seabird": electronic supplementary material S1

**Table S1**: Performance of the set of three candidate linear mixed models fitted to investigate the extent to which nest-site characteristics could predict **flight initiation distance (FID)** behaviour of the breeding pairs. Model(s) in bold were used for the estimation of parameters (ΔAICc < 2). Our set of candidate models was composed of models that included the number of FID repetitions (Repetition), the starting distance of the FID (Start), the area more used by humans (Area), elevation of the nesting site (Elevation) and density of Common gull nests within a 5-meter radius (Density) and the two-way interactions between Elevation and Density. The breeding pair identity was included as a random effect in each model to account for the non-independence of data originating from the same pair and avoid pseudo-replication issues, and the year the data were collected to control for unexplained variance due to inter-annual variation. AICc is the value of the corrected Akaike’s Information Criterion, and K is the number of estimated parameters for each model. The ranking of the models is based on the differences in the values for ΔAICc and on the Akaike weights (AICw).

| **Models** | **K** | **AICc** | **ΔAICc** | **AICw** |
| --- | --- | --- | --- | --- |
| ***Influence of nest-site characteristics*** | | | | |
| **Repetition + Start + Area + Density + Elevation** | **9** | **6164.4** | **0.00** | **0.60** |
| ***Influence of nest-site characteristics and their interaction*** | | | | |
| **Repetition + Start + Area + Density + Elevation +**  **Density*Elevation** | **10** | **6165.3** | **0.84** | **0.40** |
| ***No influence of nest-site characteristics*** | | | | |
| Repetition + Start + Area | 7 | 6188.9 | 24.44 | 0.00 |

**Table S2**: Performance of the set of three candidate generalised linear models (Binomial family) fitted to investigate the extent to which flight initiation distance (FID) behaviour is linked to **hatching success** per breeding pair. Model(s) in bold were used for the estimation of parameters (ΔAICc < 2). Our set of candidate models was composed of models that included mean adjusted FID (FID), elevation of the nesting site (Elevation), density of Common gull nests within a 5-meter radius (Density), Year of data collection (Year), and the two-way interactions between Elevation and FID and Density and FID. AICc is the value of the corrected Akaike’s Information Criterion, and K is the number of estimated parameters for each model. The ranking of the models is based on the differences in the values for ΔAICc and on the Akaike weights (AICw).

| **Models** | **K** | **AICc** | **ΔAICc** | **AICw** |
| --- | --- | --- | --- | --- |
| ***No influence of FID*** | | | | |
| **Density + Elevation + Year** | **5** | **636.5** | **0.00** | **0.50** |
| ***Influence of FID*** | | | | |
| **Density + Elevation + Year + FID** | **6** | **636.9** | **0.31** | **0.43** |
| *Influence of FID according to Density and Elevation* | | | | |
| Density + Elevation + Year + FID +  FID*Density + FID*Elevation | 8 | 640.4 | 3.83 | 0.07 |

**Table S3**: Performance of the set of three candidate generalised linear models (Binomial family) fitted to investigate the extent to which flight initiation distance (FID) behaviour is linked to the **probability of having at least 1 chick** per breeding pair. Model(s) in bold were used for the estimation of parameters (ΔAICc < 2). Our set of candidate models was composed of models that included mean adjusted FID (FID), elevation of the nesting site (Elevation), density of Common gull nests within a 5-meter radius (Density), Year of data collection (Year), and the two-way interactions between Elevation and FID and Density and FID. AICc is the value of the corrected Akaike’s Information Criterion, and K is the number of estimated parameters for each model. The ranking of the models is based on the differences in the values for ΔAICc and on the Akaike weights (AICw).

| **Models** | **K** | **AICc** | **ΔAICc** | **AICw** |
| --- | --- | --- | --- | --- |
| ***No influence of FID*** | | | | |
| **Density + Elevation + Year** | **5** | **234.4** | **0.00** | **0.70** |
| *Influence of FID* | | | | |
| Density + Elevation + Year + FID | 6 | 236.4 | 2.02 | 0.25 |
| *Influence of FID according to Density and Elevation* | | | | |
| Density + Elevation + Year + FID +  FID*Density + FID*Elevation | 8 | 239.7 | 5.29 | 0.05 |

**Table S4**: Performance of the subset of candidate linear models within a ΔAICc < 2 fitted to investigate the extent to which flight initiation distance (FID) behaviour is linked to the **clutch mass** per breeding pair. Model(s) in bold were used for the estimation of parameters (ΔAICc < 2). Our set of candidate models was composed of models that included mean adjusted FID (FID), elevation of the nesting site (Elevation), density of Common gull nests within a 5-meter radius (Density), Year of data collection (Year), Number of eggs in the nest (Eggs), and the two-way interactions between Elevation and FID and Density and FID. AICc is the value of the corrected Akaike’s Information Criterion, and K is the number of estimated parameters for each model. The ranking of the models is based on the differences in the values for ΔAICc and on the Akaike weights (AICw).

| **Models** | **K** | **AICc** | **ΔAICc** | **AICw** |
| --- | --- | --- | --- | --- |
| ***No influence of FID*** | | | | |
| **Density + Elevation + Year + Eggs** | **9** | **1829.7** | **0.00** | **0.70** |
| ***Influence of FID*** | | | | |
| **Density + Elevation + Year + Eggs + FID** | **10** | **1831.6** | **1.90** | **0.27** |
| *Influence of FID according to Density and Elevation* | | | | |
| Density + Elevation + Year + Eggs + FID +  FID*Density + FID*Elevation | 12 | 1836.0 | 6.31 | 0.03 |

**Table S5**: Performance of the subset of candidate linear models within a ΔAICc < 2 fitted to investigate the extent to which flight initiation distance (FID) behaviour is linked to the **brood mass** per breeding pair. Model(s) in bold were used for the estimation of parameters (ΔAICc < 2). Our set of candidate models was composed of models that included mean adjusted FID (FID), elevation of the nesting site (Elevation), density of Common gull nests within a 5-meter radius (Density), Year of data collection (Year), Number of hatchlings in the nest (Hatchlings), and the two-way interactions between Elevation and FID and Density and FID. AICc is the value of the corrected Akaike’s Information Criterion, and K is the number of estimated parameters for each model. The ranking of the models is based on the differences in the values for ΔAICc and on the Akaike weights (AICw).

| **Models** | **K** | **AICc** | **ΔAICc** | **AICw** |
| --- | --- | --- | --- | --- |
| ***No influence of FID*** | | | | |
| **Density + Elevation + Year + Hatchlings** | **8** | **1488.5** | **0.00** | **0.63** |
| ***Influence of FID*** | | | | |
| **Density + Elevation + Year + Hatchlings + FID** | **9** | **1490.2** | **1.67** | **0.27** |
| *Influence of FID according to Density and Elevation* | | | | |
| Density + Elevation + Year + Hatchlings + FID +  FID*Density + FID*Elevation | 11 | 1492.1 | 3.62 | 0.10 |

**Table S6**: Performance of the subset of candidate linear models within a ΔAICc < 2 fitted to investigate the extent to which flight initiation distance (FID) behaviour is linked to the **laying date of the 1st egg** per breeding pair. Model(s) in bold were used for the estimation of parameters (ΔAICc < 2). Our set of candidate models was composed of models that included mean adjusted FID (FID), elevation of the nesting site (Elevation), density of Common gull nests within a 5-meter radius (Density), Year of data collection (Year), and the two-way interactions between Elevation and FID and Density and FID. AICc is the value of the corrected Akaike’s Information Criterion, and K is the number of estimated parameters for each model. The ranking of the models is based on the differences in the values for ΔAICc and on the Akaike weights (AICw).

| **Models** | **K** | **AICc** | **ΔAICc** | **AICw** |
| --- | --- | --- | --- | --- |
| ***Influence of FID according to Density and Elevation*** | | | | |
| **Density + Elevation + Year + FID +**  **FID*Density + FID*Elevation** | **9** | **1121.4** | **0.00** | **0.59** |
| ***Influence of FID*** | | | | |
| **Density + Elevation + Year + FID** | **7** | **1122.5** | **1.14** | **0.34** |
| *No influence of FID* | | | | |
| Density + Elevation + Year | 6 | 1125.6 | 4.17 | 0.07 |

**Table S7**: Performance of the subset of candidate linear mixed models within a ΔAICc < 2 fitted to investigate the extent to which flight initiation distance (FID) behaviour is linked to the **egg mass at laying** per breeding pair. Model(s) in bold were used for the estimation of parameters (ΔAICc < 2). Our set of candidate models was composed of models that included mean adjusted FID (FID), elevation of the nesting site (Elevation), density of Common gull nests within a 5-meter radius (Density), Year of data collection (Year), Laying rank of the egg (EggNb) and the two-way interactions between Elevation and FID and Density and FID. Pair ID and Year were included as random intercept terms to account for the non-independence of data. AICc is the value of the corrected Akaike’s Information Criterion, and K is the number of estimated parameters for each model. The ranking of the models is based on the differences in the values for ΔAICc and on the Akaike weights (AICw).

| **Models** | **K** | **AICc** | **ΔAICc** | **AICw** |
| --- | --- | --- | --- | --- |
| ***No influence of FID*** | | | | |
| **Density + Elevation + Year + EggNb** | **8** | **3406.1** | **0.00** | **0.67** |
| ***Influence of FID*** | | | | |
| **Density + Elevation + Year + EggNb + FID** | **9** | **3407.8** | **1.67** | **0.29** |
| *Influence of FID according to Density and Elevation* | | | | |
| Density + Elevation + Year + EggNb + FID +  FID*Density + FID*Elevation | 11 | 3411.6 | 5.52 | 0.04 |

**Table S8**: Performance of the subset of candidate linear mixed models within a ΔAICc < 2 fitted to investigate the extent to which flight initiation distance (FID) behaviour is linked to the **hatchling mass** per breeding pair. Model(s) in bold were used for the estimation of parameters (ΔAICc < 2). Our set of candidate models was composed of models that included mean adjusted FID (FID), elevation of the nesting site (Elevation), density of Common gull nests within a 5-meter radius (Density), Year of data collection (Year), Laying rank of the egg the chick hatched from (EggNb) and the two-way interactions between Elevation and FID and Density and FID. Pair ID and Year were included as random intercept terms to account for the non-independence of data. AICc is the value of the corrected Akaike’s Information Criterion, and K is the number of estimated parameters for each model. The ranking of the models is based on the differences in the values for ΔAICc and on the Akaike weights (AICw).

| **Models** | **K** | **AICc** | **ΔAICc** | **AICw** |
| --- | --- | --- | --- | --- |
| ***No influence of FID*** | | | | |
| **Density + Elevation + Year + EggNb** | **8** | **2355.4** | **0.00** | **0.43** |
| ***Influence of FID according to Density and Elevation*** | | | | |
| **Density + Elevation + Year + EggNb + FID +**  **FID*Density + FID*Elevation** | **11** | **2355.6** | **0.25** | **0.38** |
| ***Influence of FID*** | | | | |
| **Density + Elevation + Year + EggNb + FID** | **9** | **2356.9** | **1.55** | **0.20** |
